# Pinetree: a step-wise gene expression simulator with codon-specific translation rates

**DOI:** 10.1101/339994

**Authors:** Benjamin R. Jack, Claus O. Wilke

## Abstract

**Motivation:** Stochastic gene expression simulations often assume steady-state transcript levels, or they model transcription in more detail than translation. Moreover, they lack accessible programming interfaces, which limits their utility.

**Results:** We present Pinetree, a step-wise gene expression simulator with codon-specific translation rates. Pinetree models both transcription and translation in a stochastic framework with individual polymerase and ribosome-level detail. Written in C++ with a Python front-end, any user familiar with Python can specify a genome and simulate gene expression. Pinetree was designed to be efficient and scale to simulate large plasmids or viral genomes.

**Availability:** Pinetree is available on GitHub (https://github.com/benjaminjack/pinetree) and the Python Package Index (https://pypi.org/project/pinetree/).

## 1 Introduction

Advances in synthetic biology enable biologists to precisely manipulate gene expression. As the complexity of genetic engineering techniques increases, biologists need tools to simulate and predict gene expression. Computational models of gene expression fall into three broad categories based on their granularity and underlying assumptions. Differential equation-based models treat the gene as the smallest simulated unit [1–3]. Totally asymmetric simple exclusionary process (TASEP) models account for the step-wise motion of individual ribosomes on transcripts, and provide mean-field approximations of protein production [4–6]. Stochastic simulations offer the most detailed tracking of individual molecules. However, existing stochastic models support either step-wise transcription or step-wise translation, but not both [7, 8]. The simulator we present here, Pinetree, extends these prior models. Pinetree tracks the stochastic movements of individual molecules in both transcription and translation.

## 2 Results

Each Pinetree simulation begins with a single copy of a genome, and a pool of free polymerases and ribosomes (Fig. 1A). The simulation proceeds according to the Gillespie algorithm [9] in discrete, dynamic time steps. RNA polymerases bind to promoters in the genome, initiate transcription, and generate individual transcripts by transcribing one base pair per time step (Fig. 1B). As a given transcript extends from a polymerase, ribosomes bind to exposed ribosome binding sites and initiate translation. These ribosomes translate one base per time step, and, depending on the codon, may translate at different speeds. Ribosomes may collide with and stall behind upstream ribosomes or the RNA polymerase synthesizing the transcript on which the ribosome is translating. Likewise, polymerases may collide with and stall behind upstream polymerases.

**Figure 1:**
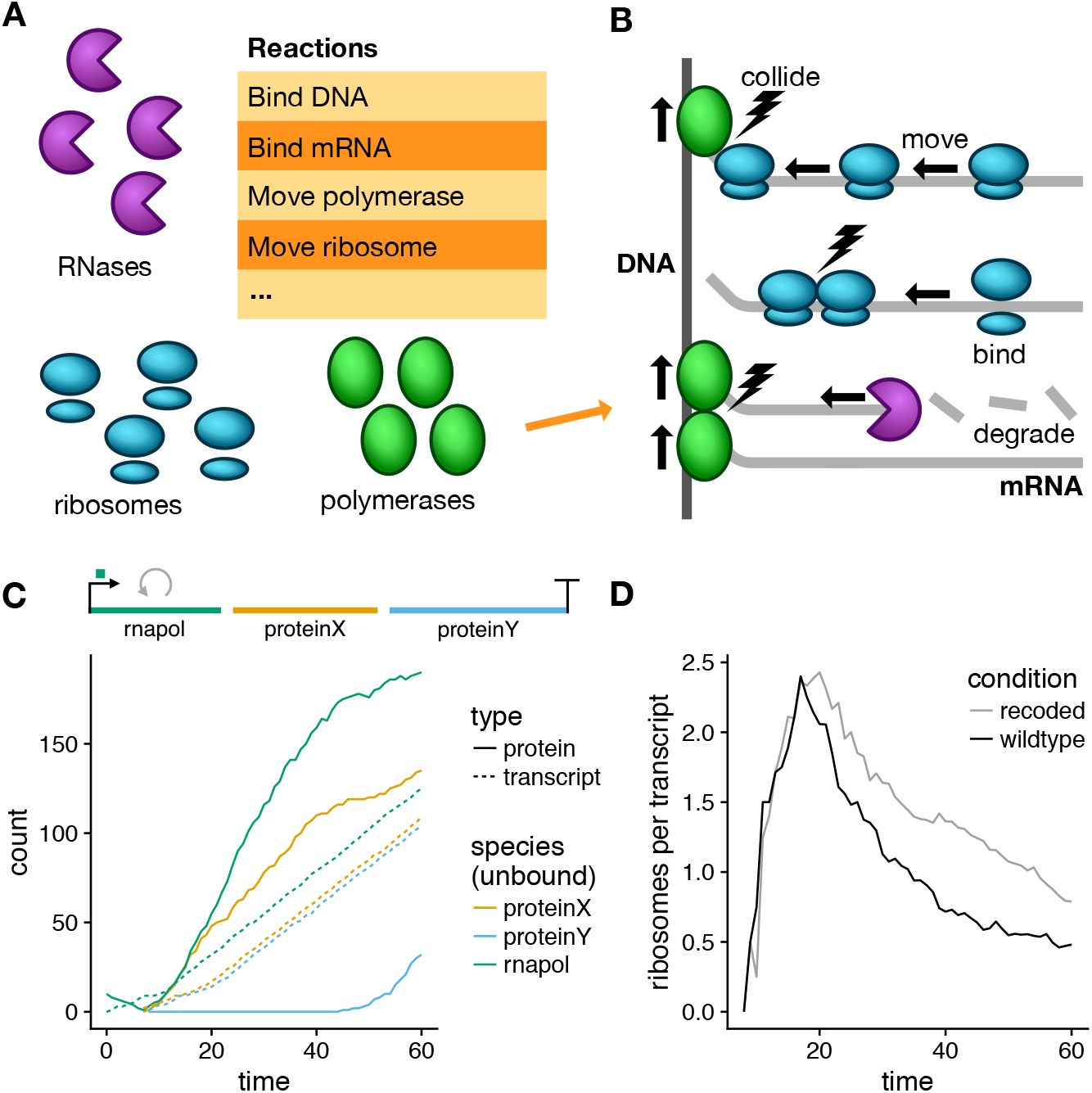
Design and example output of Pintree. Pinetree is a generic gene expression simulator that tracks individual RNA polymerases on DNA, and ribosome and RNases on mRNA transcripts. (A) A specialized set of reactions mediates binding and movement of ribosomes, RNA polymerases, and RNases. These reactions convert molecules from a species pool into individually modeled ribosomes, polymerases, and RNases. (B) Pinetree tracks individual ribosomes and RNA polymerases on transcripts and DNA, respectively. Lightning bolts represent potential collisions. Ribosomes may begin translation before a full transcript has been synthesized by the RNA polymerases. Likewise, an RNase may begin degrading an RNA transcript before the polymerase has synthesized the full transcript. Together, Pinetree allows the end user to define arbitrary reactions between molecules in the free species pool, while still modeling transcription and translation at the single-molecule level. (C) Simulation of a three-gene plasmid with one gene (proteinY) recoded to use rare codons. No RNases are present in this simulation, so no degradation occurs. Translation of proteinY is slower than that of proteinX and rnapol. (D) Ribosome densities on the transcript of proteinY are higher in the recoded gene than in the wildtype. The recoded gene has lower per-codon translation rates than that of the wild type.

To test the design and implementation of Pinetree, we developed a three-gene plasmid simulation. The plasmid was simulated in a mini-cell containing 100 ribosomes, and 10 RNA polymerases. We assumed stable transcripts that do not degrade over the course of the simulation so the system never reached equilibrium. Our goal was to determine if changes in codon usage bias produced changes in protein abundance that are consistent with our current understanding of molecular biology. The simulated three-gene plasmid contained a promoter activated by an RNA polymerase immediately upstream of the polymerase gene. Thus, expression of the RNA polymerase was autoregulatory. There was only a single terminator, so all transcripts were polycistronic. We constructed two versions of this plasmid. The first contained uniform translation rates. In the second, all preferred codons in the third gene (Y) were replaced with rare codons. Since all three genes are controlled by the same promoter, protein abundance in this model is determined by the order of the genes, the length of each gene, and codon usage (Fig. 1C). As expected, we observed a decrease in protein abundance for gene Y in the recoded plasmid but no difference in transcript abundances between the two plasmids (not shown). Along gene Y, we also observed a higher rate of ribosome collisions and stalling and an increase in ribosomal density in the recoded plasmid (Fig. 1D).

To test the performance of Pinetree, we simulated the infection of a simplified E. coli cell model by bacteriophage T7. The T7 genome is 40kb and contains 60 genes, with an infection cycle of 12 minutes. Simulating gene expression of this phage across the infection cycle took approximately 3 hours.

## 3 Discussion

Pinetree offers two main advantages over prior simulations of translation. First, Pinetree simulates dynamic transcript abundances and transcript lengths. Transcript abundances can change over the course of the simulation, and precise transcript lengths can be determined dynamically through mechanisms like terminator readthrough. Past simulations require that you pre-specify both transcript abundances and transcript lengths [4–7,10]. Second, unlike TASEP models, ribosome movements are modeled explicitly on each transcript [4–6]. Together, these two properties of Pinetree allow the user to explore non-steady state dynamics of gene expression at the single base pair resolution, all without pre-specifying transcript lengths or abundances.

Of course, such a detailed simulation structure comes at a performance cost. While implemented in C++ and designed to be highly-efficient, Pinetree’s performance will never be able to compete with simulations in which translation rates are estimated via mean-field approximations, and simulations that do not model transcription. Future versions of Pinetree could implement alternate versions of the Gillespie algorithm that make some approximations in order to decrease run times and increase efficiency [11, 12].

Despite the performance cost of single-base pair resolution, we have demonstrated that Pinetree is scalable to the size of viral genomes. We envision that Pinetree could be used to simulate small bacterial genomes, and facilitate *in silico* viral evolution experiments.

## 4 Methods

We designed designed and prototyped Pinetree in Python. After validating the Python implementation, we rewrote portions of the Python source code into C++ to improve performance. A C++ header library pybind11 provides Python wrappers for C++ classes so that C++ objects interact seamlessly with the Python interpreter.

Pinetree is available as a user-friendly Python package. All Pinetree source code is available on Github (https://github.com/benjaminjack/pinetree) and the Python Package Index (https://pypi.org/project/pinetree/). All documentation is available on Read the Docs (https://pinetree.readthedocs.io/). We are in the process of archiving Pinetree version 0.1.0 with documentation on Zenodo.

